# Neuromorphic Model of Hippocampus Place cells using an Oscillatory Interference Technique for Hardware Implementation

**DOI:** 10.1101/2022.07.18.500440

**Authors:** Zhaoqi Chen, Alia Nasrallah, Milad Alemohammad, Masanori Furuta, Ralph Etienne-Cummings

**Affiliations:** The Johns Hopkins University, Baltimore, Maryland USA; Toshiba Corporation, Kawasaki, Japan

**Keywords:** Oscillatory Interference, Place cells, SLAM, Neuromorphic Engineering, Hardware Implementation

## Abstract

In this paper, we propose a simplified and robust model for Place cell generation based on the Oscillatory Interference (OI) model concept. Aiming toward hardware implementation in bio-inspired Simultaneous Localization and Mapping (SLAM) systems for mobile robotics, we base our model on logic operations that reduce its computational complexity. The model compensates for parameter variations in the behaviors of the population of constituent Theta cells, and allows the Theta cells to have square-wave oscillation profiles. The robustness of the model, with respect to mismatch in the Theta cell’s base oscillation frequency and gain—as a function of modulatory inputs—is demonstrated. Place cell composed of 48 Theta cells with base frequency variations with a 25% standard deviation from the mean and a gain error with 20% standard deviation from the mean only result in a 20% deformations within the place field and 0.24% outer side lobes, and an overall pattern with 0.0015 mean squared error on average. We also present how the model can be used to achieve the localization and path-tracking functionalities of SLAM.

## 1. Introduction

Most animals, including humans, have the capability to navigate through complex environments. They can keep track of their locations relative to their homes, and can plan and memorize paths through clutter as they advance towards their targets. When exploring unfamiliar environments, animals can also store memories that map their travel, locate scents and visual cues and, ultimately, assign meaning to various aspects of the environment. Such capability is of interest to the robotics community and is known as Simultaneous Localization and Mapping, or SLAM, which is a computational problem aimed at constructing a map for an unknown environment and localizing the robot in that environment while exploring it [1].

SLAM has long attracted interest in mobile robotics, and typically gets inputs from heavily over-sensored robots. It often uses complex mathematical algorithms, requiring in high computation rates from high energy consuming systems. However, animals can achieve high-performance SLAM without, to our understanding, the ability to perform any overt mathematical computation [2-6]. Neuroscience researchers are attempting to explain how the nervous system encodes the spatial environment in which we navigate, and how that contributes to path planning and navigation [7-11][20-24]. We believe that neuroscience research can inspire more efficient SLAM implementations than traditional approaches emerging from the robotics community. The key metric for efficiency will be compactness of the models, simplicity of the computations, and ease of hardware implementation.

Many nerve cells located in the hippocampus have been found to encode location or features in the environment and participate in navigation [2-5]. Distinguished by their behaviors, two types of such cells were termed ‘Grid cells’[2] and ‘Place cells’ [3]. In particular, in 1971, O’Keefe discovered Place cells in the hippocampus of the rat, which responded only when the animal is within a particular spatial location, defined as place field [2]. In 2005, Moser found Grid cells that are activated periodically and regularly as the animal explores a given space [3]. These results were significant enough to warrant a Nobel Prize in 2014. As animals explore unfamiliar environments, new location-specific cells and maps are generated and remembered. Grid and place neurons are believed to play a critical role in spatial encoding and subsequent, localization and navigation functionalities [4-6].

To explain how the Place and Grid cells may emerge from the known oscillatory functions of neurons in the hippocampus, the Oscillatory Interference (OI) model was introduced by Burgess et al. [7]. It attributes the signal of the Place and Grid cells to summations of lower-level neurons, named ‘Theta cells’, that have specific frequency tunings to head direction and speed of movement in space. This was further supported by Welday et al.’s discovery of the Theta cells’ frequency response to the animal’s travel speed and direction [8]. The model presented in this paper will leverage these results because of their elegance and ease of implementation in hardware.

Another class of model used to explain the formation of Place and Grid cells is the Continuous Attractor Networks (CAN) model which generates Grid cell activity based on a collecion of cells that projects and receives unsymmetric inhibition connections with their surrounding neighbors, as well as a distinct directional tuned inputs for each of those cells[9]–[11].

Enlightened by these findings, many biologically inspired SLAM algorithms, such as NeuroSLAM or RatSLAM, offer methods that operate with limited small number of sensors and consume low power [12]–[16]. NeuroSLAM promises performance that can mimic living organisms, with the potential to surpass the performance of current computational approaches. While previous neural inspired SLAM implementations take inspiration from general behavior models of spatial encoding and navigation in rat hippocampus and para-hippocampus regions, their implementation, which is mainly in hardware, still employs complex computational techniques [17]. Hence, by failing to model the elegancy of lower-level neural circuitries found in the hippocampus, these models are typically cumbersome and cannot take advantage of the computational and power consumption efficiency offered by neuromorphic systems, in both software and hardware incarnations.

One reason that the traditional SLAM approach has shied away from the neuromorphic approach is because the latter may require a large and complex network of neurons [10]. However, the neuromorphic approach, particularly when implemented in hardware, can take advantage of abstractions that will result in smaller hardware complexity. Consequently, we follow a philosophy of abstracted neuromorphism in this work, rather than of direct neural mimicry.

As an example of the type of abstracted neuromorphic approach that we have taken, we can consider the method for implementing the Theta cells, which are the velocity-sensitive neurons found in the hippocampus. These Theta cells, which are required by the OI model to generate Grid cells [7], are assumed to have a base oscillatory frequency that varies according to a cosine tuning curve that scales according to the dot-product between the input velocity vector and the preferred velocity vector of the cell, as indicated in equation (1). In Welday et al.’s proposed model for Place cell formation [8], a uniform gain for the frequency change as a function of travel velocity, and a uniform base frequency, are necessary when interfering independent oscillators with distinct preferred velocities. However, it is unrealistic to expect biological neurons to have these uniform behaviors. Similarly, it is impossible for different circuit modules implementing Theta cells, particularly when implementated with analog circuit, to have the same base frequency and vary in the same way when the inputs are varied. Hence, any model that uses hardware neurons or oscillators must provide a pathway to reduce the impact of these non-uniformity amoung various Theta cells.

The CAN model offers its own implementation challenges. In the CAN model, a collection of oscillating cells’ frequencies are regulated by input velocity and their weighted inhibitory connections with its surrounding cells, making the interconnections very complex for hardware implementation. Furthermore, as a population based model, scaling down the number of such cells to reduce the complexity would reduce its robustness or even malfunction, which places a burden on system complexity.

In this paper, we propose a mathematical model for forming hippocampal Grid cells and Place cells using an Oscillatory Interference (OI) model similar to that of Welday et al. [8]. As a key distinction from Welday et al.’s model, our model interferes Theta cells using *multiplicative*, rather than *additive*, interactions. Furthermore, our model does not require the Theta cells’ base frequencies and velocity tuning gains to be identical across the interfering population. The tolerance of variation across the model’s population allows the model to better represent the realities of biological Theta cells, and to be more robust to the non-idealities of implementations with analog (or mixed-signal) hardware, which are typically much more space and power efficient than their digital counterparts. We then demonstrate the improved capability of our model to instantiate a Place cell at any location, despite the expected variation and mismatching between the Theta cells. Lastly, we show how the model can be used for path tracking and how to deal with the accumulation of position error as the agent (or animal) moves through space. The latter is a well know problem with “dead-reckoning” based path tracking [18]. We also demonstrate how the original model of Welday et al. behaves with the same levels of variations and errors in the Theta cells. Lastly, we demonstrate the robustness of the model as a function of model complexity (i.e. number of oscillators per Place cell), oscillator variations in both base frequency and its response gain as a function of travel velocity.

This paper is organized as follows: first, in section 2, we briefly introduce the prevailing models for spatial cell formation. In section 3, we demonstrate the models through simulation, and introduce our multiplicative construction method for neuromorphic Place cells. In section 4, we characterize the model by inducing variations among Theta cell behaviours. In section 5, we improve our model to accommodate the variation and reduce the variation’s impact on spatial accuracy of the Place cells. In section 6, we propose a path-tracking mechanism incorporating our Place cell model. In section 7, we conduct simulations to examine the performance of the Place cell model under various conditions and to demonstrate path-tracking functionality. Finally, we conclude this paper by discussing future directions of this work.

## 2. Related models

As mentioned previously, there exist two major theories on how place and Grid cells generate their response to an animal’s location. The first model, the Oscillatory Interference (OI) model proposed by Burgess et al. concisely describes which Grid cell patterns are formed with a summation of distinct Theta cells’ signals, with each Theta cell having a distinctive preferred direction. Individual Theta cells oscillate at a frequency that scales linearly with the dot-product between its preferred velocity and the travel velocity. The OI model has a nice modular structure in which each Grid cell is independent, and the operation for creating Grid cells requires only summation and thresholding. Furthermore, the model has a “Fourier” construction nature, which allows the Place and Grid cells to be more compact when more base Theta cells are introduced in the summation. However, the model expects each Theta cell to have the same base frequency of operation, e.g. 8Hz, and is susceptible to mismatch between Theta cells’ behaviors. When the Theta cells have different base frequencies and/or different tuning gains, the model does not produce compact Place and Grid cells, and they may have multiple response lobes. However, Welday’s discovery [8] that biological Theta cells vary in their frequencies, either idling or under movement, is not surprising. Hence, if Theta cells are implemented as ring oscillators as they suggested, then it would be difficult to maintain an array of independent ring oscillators with both a same oscillation frequency as well as individual frequency response curves.

The other prevailing model is the Continuous Attractor Network (CAN) model. It is a population based model where each cell needs the interconnection around its neighbors to operate. In the CAN model, each cell has a unsymmetric inhibition connection to its neighbors based on a prefered direction. Each cell also receives an input that is stronger when the subject is traveling along the prefered direction. Thus when the subject is traveling along that preferred direction, the cell will become more active then push the pattern towards that direction through its unsymmetric inhibition projecting to the neighboring cells. The model maintains stability through the population that has various preferred direction that cancel out the asymmetry and exhibits a equilibrium pattern overall. The downside is also obvious since it needs a population to function which is not a modular design. And a large number of carefully weighted inhibition interconnection is also needed to form such a network, making it less attractive to a neuromorphic implementation.

Literature suggests that Grid cells plays a crucial role for formation of Place cells to achieve path integration when visual cues are absent [19,20]. Solstad et al. [21] and Blair et al. [22] proposed models that form Place cells by summing sophisticatedly weighted Grid cells, suggesting a hierarchical structure from Theta cells to Grid cells then to Place cells. However, such weighting imposes another burden on circuit design, especially when a large number of Place cells are needed to tessellate a space. Hence, a simplified Place cell construction approach is proposed in this paper. In addition, we also discuss a explanation for the necessity of such hierarchy.

## 3. Oscillatory Interference models under ideal conditions

In order to preserve simplicity, our model takes the OI model as its fundamental structure. As predicted [7], [23]– [26] and later discovered by Welday et al. [8], the basic component of the OI model originates from the Theta cells signals that act like velocity-controlled oscillators. Each Theta cell has an oscillating frequency that is modulated by the animal’s movement direction and speed, as shown in equation (1), in which the firing phase accumulation is synchronized with displacement along a preferred direction.

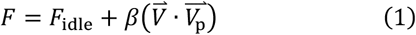

Here *F* denotes the oscillation frequency of the Theta cell, which depends on the dot-product between its preferred velocity 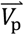 and the animal’s travel velocity 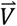. *β* is the frequency response factor or gain, which relates the traveling speed with oscillation frequency, with units of *Hz/unit speed*. The idling frequency *F*_*idle*_ denotes the frequency when the input velocity is zero. Welday et al. experimentally observed that Theta cells have various preferred velocity vectors, and believe that they generate the fundamental signals for spatial encoding through oscillatory interference.

### 3.1 Oscillatory Interference to form Grid cells

The core concept of the OI model is to form Place cells and Grid cells simply by summing one Theta cell’s velocity-regulated oscillation and another reference oscillator with the idling frequency *F*_*idle*_, using the standard trigonometric identity below:

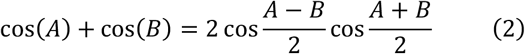

The cosine containing the difference between the two frequencies would represent one Theta cell’s deviation from either a reference cell or another Theta cell. Burgess et al. [7] suggested that forming Grid cells is possible, and indeed concise, by interference between merely a few Theta cells. A one-dimensional Grid cell with a periodic response to traveling in the Theta cell’s preferred direction can be formed by just interfering that Theta cell with an idling frequency, or, equivalently, with a Theta cell with 0-valued preferred velocity, resulting in the interference pattern shown in figure 1 and its signal below:

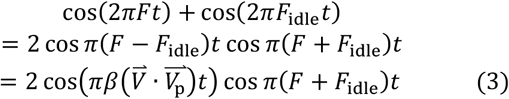

Each point on the spatial firing pattern figure is calculated by traveling along a direct path from the origin to that point with a pre-set fixed speed 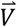, then plotting the instantaneous signal value at that location through equation (3). The vertical gratings signify that this Grid cell will fire periodically while the subject is traveling in its preferred direction, as the gratings will always be perpendicular to the preferred velocity vector. According to equation (3), the resulting signal will have a low-frequency component that depends only on the Theta cell’s frequency response to an input velocity. After forming the grating, it is straightforward to form the hexagonal Grid cells by interfering with an additional Theta cell which has a preferred direction, say 60° apart, to form an interference pattern that resembles the response of Grid cells, shown in figure 1.

**Figure 1:**
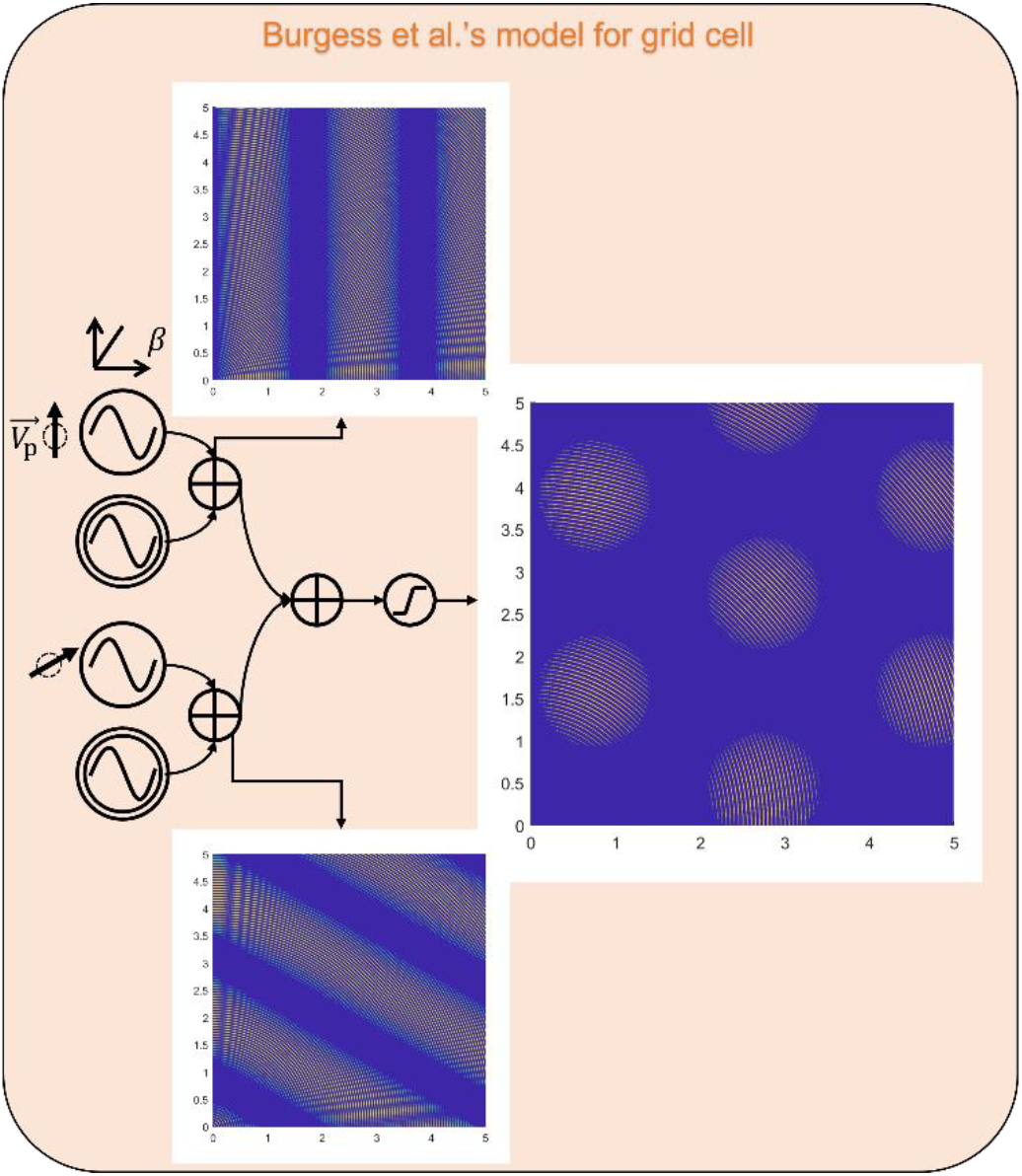
a 2-D Grid cell by interfering two 1-D Grid cells. Each 1-D Grid cell is formed by interfering a theta cell and an idling oscillator. The theta cells have the same frequency response factor β but different preferred direction 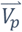. Refer Symbols to figure 2.

### 3.2 Oscillatory Interference to form Place cells

Although many researchers suggest that Place cells are formed by Grid cells, Welday et al. proposed a hypothesized method for forming Place cells directly from Theta cells. If we focus on the low-frequency component, i.e. the envelope, of the interference result in equation (3). The overall envelope of the interference result from *N* Theta cells can be described by equation (4).

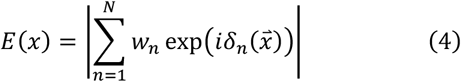

Here, *E*(*x*) is the envelope evaluated at the spatial location *x* through a vector 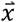 with respect to the origin, *w*_*n*_ is a weighting factor, and *i* is the imaginary unit to make use of Euler’s form for the phase *δ*_*n*_(*x*). It is the phase accumulation that deviates from the idling frequency, as mentioned in equation (3). It can be computed as equation (5), with an additional initial phase shift 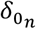 to add a degree of freedom for distinguishing 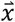.

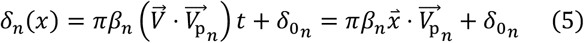

Welday et al. suggested a possible setup for Place cells. By interfering 12 Theta cells with a uniform weight of 1 and a uniform coefficient *β* but with preferred directions spaced 30° apart, a Place cell can be formed, using a properly chosen threshold, with its pattern shown in figure 2 with initial theta phases calculated from equations (4) and (5). Specifically, at the desired location *x*, set *δ*_*n*_(*x*) equals 1 then solve for 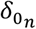 for the *n* th Theta cell’s initial phase through equation (5).

**Figure 2:**
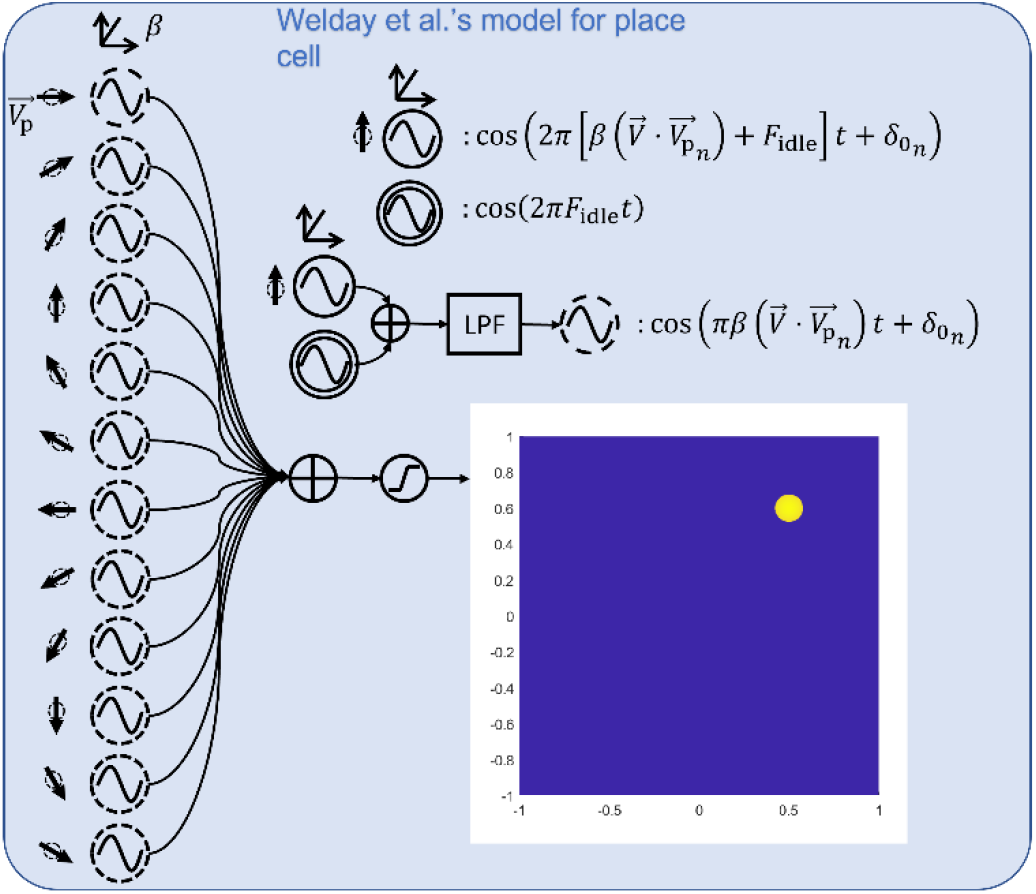
Place cell generated by Welday et al.’s model from 12 Theta cells

Though under Welday’s model, a place cell can be formed with a population of theta cells with various preferred directions other than the one shown in figure 2, issues arise when constructing such a system. First, the restrictions on the setup of preferred directions of theta cells is unclear. For example, the minimum number of directions and their orientations are not specified. Second, the nonlinear thresholding operation requires a heuristic value for the threshold that introduce ambiguity, where a threshold too high will reduce the place field quickly whereas a value too small results in a number of side lobes. The latter is not easily resolvable if the oscillations are to be implemented as square wave. Since the square wave essentially normalize all values of a sinsoid wave that is greater than zero, the interferences lined up at the peaks have no difference from those that just have positive overalappings. This losing of uniquess will cause side lobes even after thresholding, as shown in figure 3.

**Figure 3:**
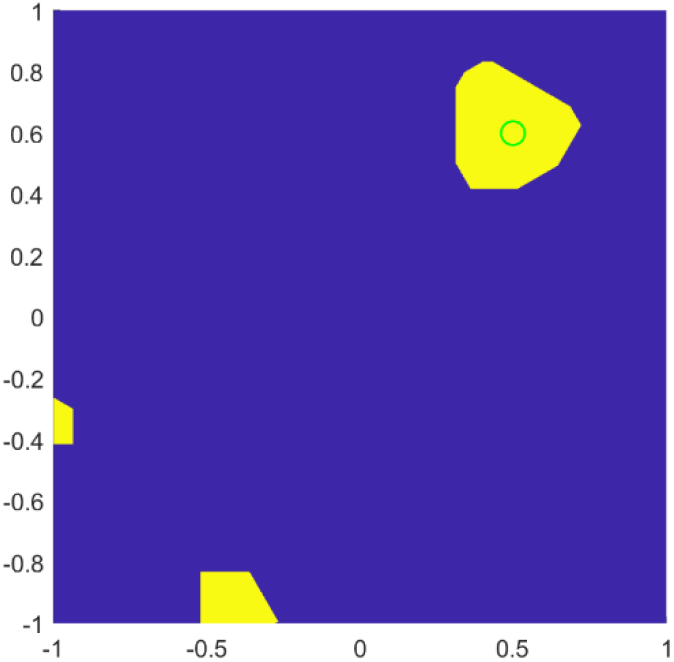
Place cell firing pattern with the same configuration from figure 2 but with square wave oscillation profile

Another important aspect of the model requires a homogenous oscillation at the idling frequency of the other speed dependent theta cells. Such synchrony might be realized in the neural circuit through shunting inhibition as suggested in [27] for the case of gamma oscillation and in [28] for pyramidal neurons both in the hippocampus CA3 where place cell resides. Shunting inhibition is a type of inhibition that regulates the membrane potential divisively rather than additively, resembles the logic AND operation.

### 3.3 Multiplicative interference and a constructive interference method for generating Place cells

To simplify hardware implementation complexity, and to remove ambiguities regarding thresholds, we adopt the logic AND operation for interference rather than addition then thresholding. Such cumulative ANDing procedure enforces constructive interference of all selected Theta cells at the location of interest, while any other side lobe will be cleared to zero by any theta envelope that is not 1. We also assume that a hardware-emulated Theta cell is likely to output square waves rather than sinusoids. In addition, quantized phase shifts are adopted here to better emulate both the neural circuitry and hardware realization of Theta cell as ring attractor [29,30] or oscillator. The AND interference operation can then be made analogous to the multiplication of two sinusoids, with the following form

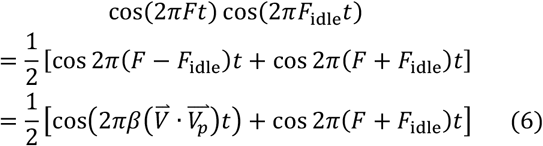

In the case of the logic AND operation, the one-half coefficient would drop off, resulting in only a superposition of the sum and difference of the frequencies, instead of modulation which made separation of the two components easier. Aside from that, the result of the envelope function is identical to that of the summation method.

To simplify the interference, using fewer Theta cells, and to reduce the complexity of phase computation, we adopt a construction method for grid- and place-cell formation inspired by the Grid cell construction method from Burgess et al. [7], which avoids the weight computation needed by the models of Solstad et al. [21] and Blair et al. [22]. Furthermore, unlike the structure proposed by Welday et al., we form Place cells from Theta cells with different frequency response factors. By introducing another degree of freedom, we can reduce the number of Theta cells needed and make the model more flexible and robust. We start by manipulating the simplest interference pattern where interference happens between one idling frequency oscillator and one Theta cell, as shown in figure 4(a); this pair acts as a one-dimensional Grid cell. Then, we form just one stripe in the field of interest by interfering gratings with the same directivity but different frequency response factors, as shown by the top and bottom rows of figure 4. This process can be viewed as summing multiple sinusoidal waves with different frequencies to form one peak. However, the Fourier analysis on either an impulse-, sinc- or Gaussian-shaped place field would result in a continuous spectrum, which is not realistically possible from a limited number of oscillations. So, we step back to redefine a Place cell as being a periodic signal with very large wavelength relative to our area of interest. According to [5], the peak separation *L* for the grating pattern can be computed with equation (7), where *β* is the frequency response factor for the Theta cell in equation (1).

**Figure 4:**
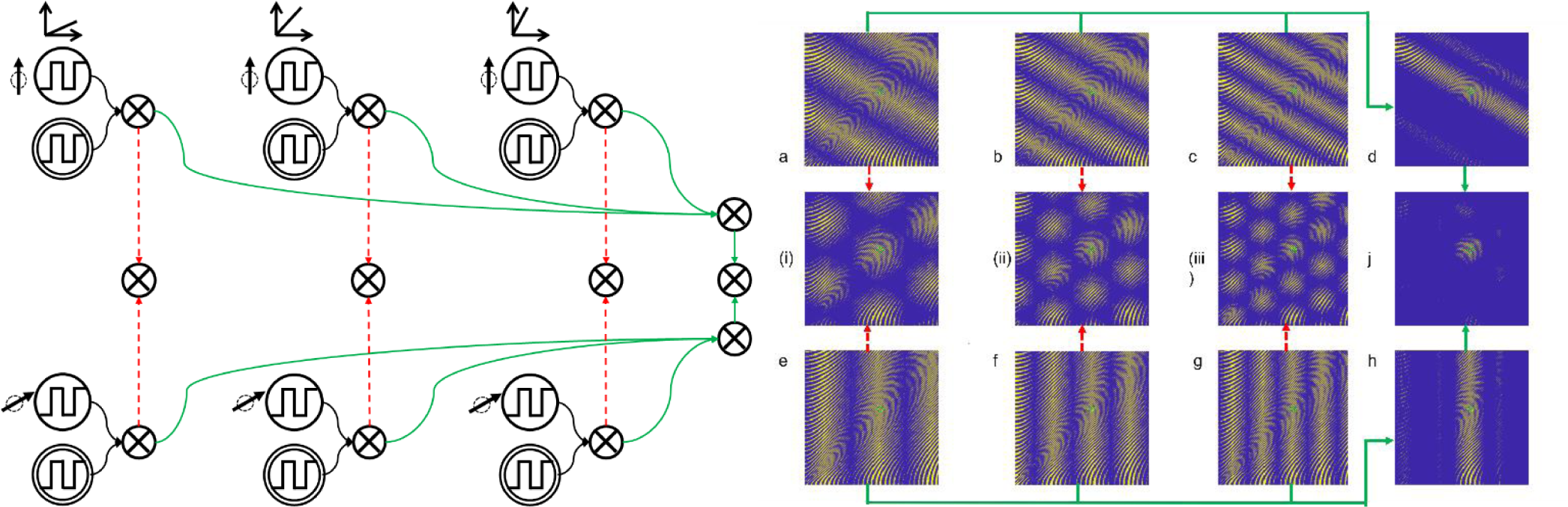
Constructive method for Place cell generation. Two groups of Theta cells with the same set of frequency response factors, but different preferred directions, are used to form the Place cell; process shown by the green arrows. Grid cells can be formed by interfere between Theta cells with same frequency response factor but different preferred directions, shown by the red arrows. And interfering (i), (ii), and (iii) would also result in j. All oscillations are square-wave and phase-shifted in 8 quantized steps to form a Place cell at the location indicated by the green circle.

**Figure 5:**
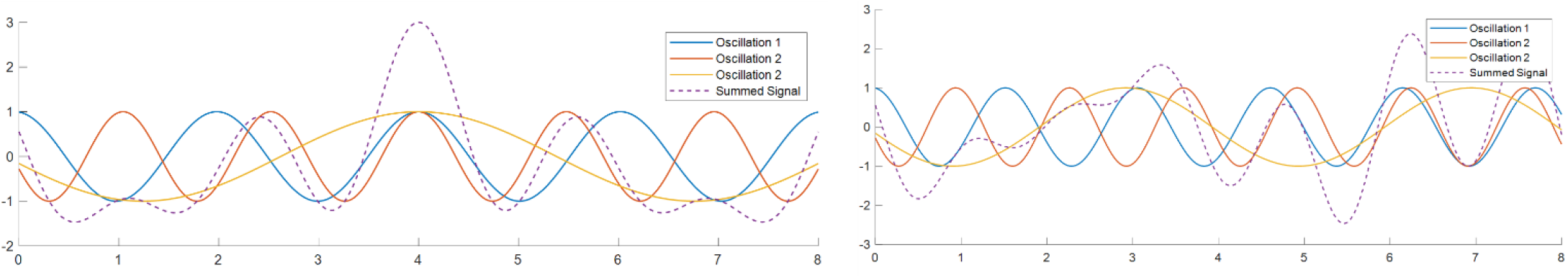
Interference signal in time domain. The left is an interference pattern of three Theta cells with initial phases calculated to constructively interfere at t = 4s under ideal condition; where the right shows a disturbed interference pattern due to the additive offset frequencies distributed from 0 to 1 Hz among the three components.

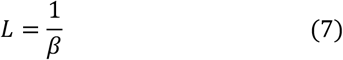

However, covering large *L* with small *β* directly is difficult in hardware implementation, and requires precise control. So instead of applying a particular value of *β* directly, we propose an interference strategy that can achieve a large value for *L* as well. Consider *β*_1_ = *pβ* and *β*_2_ = *qβ*, which results in the pattern having *p* or *q* peaks before reaching *L*. Then, the sum of the two associated signals will have their first lined-up peak at the least common multiples of *p* and *q*. Adding more such oscillators with mutually prime frequency response factors (i.e. not multiples of the others), can create gratings with very large periods, as shown in figure 4(d), (h), which are generated from three Theta cells with distinctive response coefficients *β*.

After forming a single stripe within an acceptable region, we can offset the stripes, by adding a constant phase term, in the direction of its preferred velocity vector to guarantee that a firing stripe crosses through the desired location. This is similar to constructing a surface from two orthogonal bases. To compute the phase term, we treat the desired location point, denoted as [*R, θ*] in polar coordinates, as a vector projected onto the preferred velocity vector to compute its angular frequency of oscillation along the preferred direction, [*β, θ*_*p*_]. Then, we assign one of the eight quantized phase fraction values *φ*_*i*_, which range from 0 to 7, to adjust the oscillation signal to have a peak close to the desired point, referring to equation (8). The resulting equation for the stripe pattern is illustrated in (9), where *F*_*i*_ is the frequency response of the *i*th Theta cell as defined in equation (1).

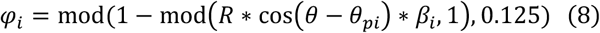

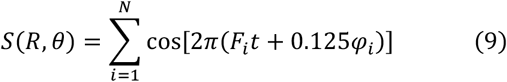

If apply the phase computation to all the oscillators used to form a single stripe in the same 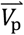 direction, we guarantee that the stripe will form along the perpendicular direction and that it passes through the desired point. Repeating this same process with further oscillators with other directivities, we intersect all of their stripes at the desired point, resulting in a single-place field as shown in figure 4(j), where we set the other group of Theta cells to have a 60° difference in their preferred velocity direction. The phase shift operation hence replaces the weight computation problem with a phase connectivity problem, or a binary weight system where only one of the eight phases of each Theta cell will be weighted 1.

As shown by the figures, the constructed stripes and Place cells have small ripples within, that reflect the high-frequency component in equation (3) and provide a potential explanation for how the phase procession exists [7]. But, for the purpose of spatial tuning, we only consider the low-frequency component, i.e. the spike envelope, which is a standard process of the OI model. In further discussions, we would filter the high-frequency component out. Thus, we implement an envelope-detection or low-pass filter stage, much like the integration process of synapses in a biological system. Eliminating the high-frequency component creates a solid Place cell that reacts to its entire place field, which is important when the number of interferences increases: as shown in figure 4, the trough of the high-frequency components of the two stripes would destructively interfere in the AND process, creating an intersection with a very small positive region. Thus, for our demonstration purposes, we define the interference operation in the frequency domain as follows:

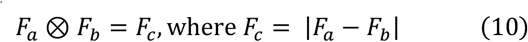

The proposed model of forming Place cells is very simple and powerful since it only needs two stripes as a basis. Moreover, there is no weighting, thresholding, or memory necessary for the interference operation, which can be implemented as combinational logic. The number of Theta cells with distinct frequency response factors needed is related to how large a region of interest is needed, since they govern the periods of the gratings. The model could be used to form any number of Place cells by applying different sets of constant phase terms to the same Theta cells’ oscillation setup. This allows us to implement the model in hardware with a rudimentary feedforward network, where a small number of voltage-controlled oscillators can serve as inputs to generate any Place cell.

Because the whole process for the OI model comprises ANDing signals, the interference for forming a single stripe for different directions can be interchanged to form Grid cells as shown in figure 4.i, ii, iii. This is an interesting discovery hinting at how the Grid cells can come into existence. But it also raises the question of whether the existence of Grid cells is required: interference could be done once at the Place cell directly, without the need for middle steps. We will, however, discover another crucial reason for why Grid cells are necessary when we start to introduce variations into the OI model, reflecting the variations inevitable in both neuromorphic hardware implementation and biological systems. We will also demonstrate that the constructive methods can even take advantage of Theta cell variations to form Place cells having unique place field with minimal side lobes over a large region.

## 4. Oscillatory Interference models under Theta cell variations

Although all the models above could generate reasonable grid- and place-cell firing patterns, they are all based on the Theta cell model described by equation (1). This model, however, is very likely to be an oversimplification. In research on Theta cells in rats [8], and when implementing Theta cells as ring oscillators in silicon, the frequency response factor *β* cannot be exactly uniform across all Theta cells. More importantly, *F*_*idle*_ also varies between Theta cells. Thus, we adjust the Theta cell equation with a constant offset frequency term, as shown in equation (11).

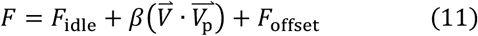

Under such variation condition, the idling frequency *F*_*idle*_ can be computed as the median of oscillation frequencies over the whole population of Theta cells under zero-velocity input. Then, *F*_offset_ can be computed for each Theta cell. This offset term does not depend on travel velocity, and significantly alters how the phase accumulation model represents space. This perspective implies—or tacitly assumes—that phase accumulation deviation happens even when the input velocity 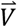 is zero. If one Theta cell interferes with another with the idling frequency as suggested by Burgess et al., the resulting frequency will be:

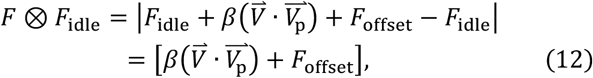

and the signal in the time domain becomes

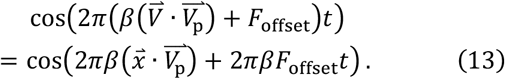

Thus, after interference involving an idling frequency, the phase accumulation is no longer solely dependent on the first term of a displacement vector 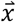 but also on the second, time-dependent, term. On a one-dimensional interference pattern graph, the phase accumulation induces curvature in the grating as shown in figure 4(a), corresponding to the greater travel time needed to reach points farther from the origin, so that the time-dependent term takes precedence. The overall frequency after interference is also adjusted by *F*_offset_ and can result in a shrinkage or expansion of the inter-peak distance *L*, resulting in the deformation of both its geometric structure and scale as shown in figure 6.

**Figure 6:**
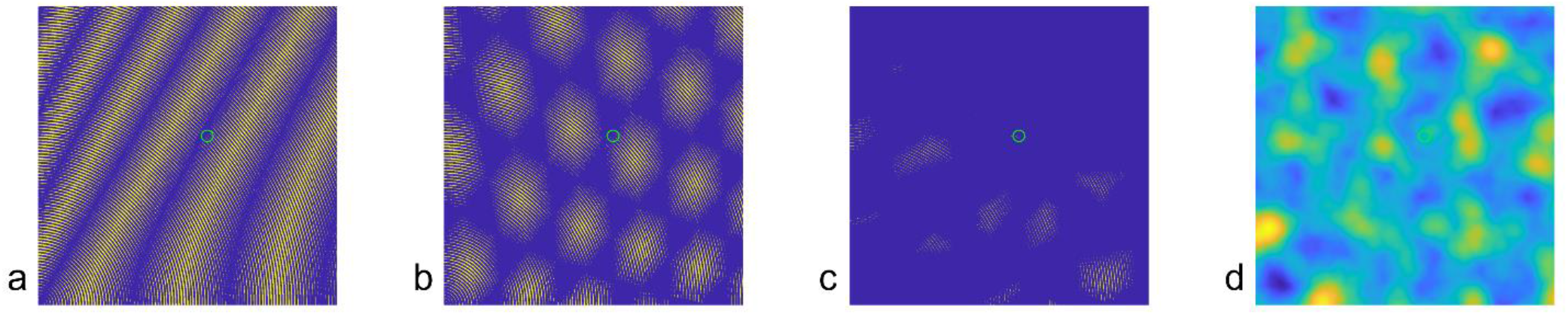
Altered firing patterns due to additively offset frequencies with a mean of 0 and a standard deviation of 0.5 Hz to the Theta cells. (a) a 1-D Grid cell pattern with from fig. 4(f) skewed by the additive offset frequency. (b) a 2-D Grid cell is generated after interfering with another Theta cell with preferred direction of 90° but also skewed by the additive offset frequency. (c) the Place cell that would be generated, as in figure 4(j). (d) the Place cell using the Welday et al.’s model using the same frequency setup as figure 2 but disturbed by the offset frequencies among the constituent Theta cells, thresholding process removed since no points is above the original threshold.

Such patterns and the equation imply that the OI model under the influence of an offset idling frequency will not provide a unique pattern that depends only on the spatial location through velocity integration. Differing routes and varying speeds during travel could disturb the phase accumulation through the time-dependent term, resulting in a completely different set of phase inputs for the Place cell, so the Place cell would not respond to its designated location. Moreover, the variations in the frequency response factor *β* even prevent the Place cell generation model proposed by Welday et al., where a set of Theta cells with uniform *β* is needed. Furthermore, as shown in figure 6(d), merely introducing the offset frequency compromises the formation of the Place cell.

We do not consider such variation merely an artifact of analog hardware implementation: achieving a totally uniform idling frequency across all Theta cells, yet retaining their ability to change frequency independently, is not consistent with biological behavior either. So, we decided that an improved version of the OI model for forming Grid cells and Place cells is critical to understanding the nature of hippocampus spatial encoding, and critical also for feasible and robust implementation in hardware. Along with the development of this improved model, we present a potential function for the Grid cells below.

## 5. Improved OI model under Theta cell variations

The variations among the frequency response factors *β*_*i*_ do not affect our proposed construction method for Place cells. In the ideal scenario, we need to deliberately construct a set of mutually prime *β*_*i*_ to form a unique stripe in the region of interest in figure 4. Under the condition of variations, the different values of the *β*_*i*_ naturally construct a large lowest common multiple that results in a large *L*, covering a large distance. In other words, our proposal does not require the capability to program the frequency response factors, as long as the actual *β*_*i*_ can be characterized. We believe that this property simplifies the Place cell model and removes substantial burden from hardware design, which we illustrate with a simulated scenario in the results section below.

The additional offset for idling frequencies across Theta cells for interference poses a more significant challenge to the principal idea of the OI model, with respect to achieving path integration through phase accumulation. In our improved model, we propose the solution in two steps: first, we propose an offset-reduction strategy of Theta cells to reduce the impact of frequency offsets. Then, we modify the phase-computation equation to accommodate the remainder of the offset frequency.

Inspecting the oscillatory interference equation (3), we can observe that interference with an idling frequency signal has the effect of removing it from the oscillation, so that the resulting signal frequency depends on the product of speed and time, which is effectively the displacement, as shown in equation (10), in the ideal case when *F*_offset_ = 0. A similar approach could be applied to remove the offset frequency. Implementing that in the basic structure of the OI model would preserve its simplicity, and thus lead to simpler hardware requirements.

Our first step in removing the offset frequency is to pair oscillators with the closest idling frequency whose interference eliminates the magnitude of the offset frequency as much as possible, as shown in equation (14). However, if we just set the second oscillator to have a preferred velocity of [0, 0], it is possible to run into the circumstance that, at a certain input speed, the first oscillator’s frequency becomes equal to the second oscillator’s idling frequency, so that the superposed oscillations generate a stationary pattern that cannot represent the distance traveled. To avoid this, we set the second oscillator to have a preferred velocity vector 180° apart from the first one. In this configuration, we effectively obtain a new oscillator that operates as shown in equation (14), with the structure as in figure 7, configured so that *F*_off_ *a*_ > *F*_off_*b*_.

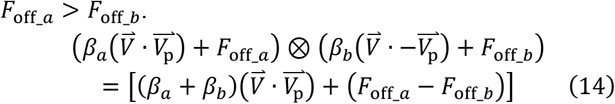

The frequency response factor is summed so to make the velocity-dependent term more predominant for the effective oscillator, while the offset frequency is suppressed. Figure 8 illustrates the suppression effect of offset-reduction between a pair of Theta cells with preferred directions 0° and 180°, and offset frequencies 0.5 Hz and 0.6 Hz, respectively. This kind of offset-reduction strategy could be repeated to obtain an acceptable offset frequency for phase computation, in terms of the number of Theta cells needed. The number of effective Theta cells would then be divided by 2^*N*^ where *N* is the number of layers. Referring to the previous discussion, the resulting effective frequency response factor (*β*_*a*_ + *β*_*b*_) can be of any value, and contributes to the feasibility of the pairing process in the framework of our construction method for place-cell generation.

**Figure 7:**
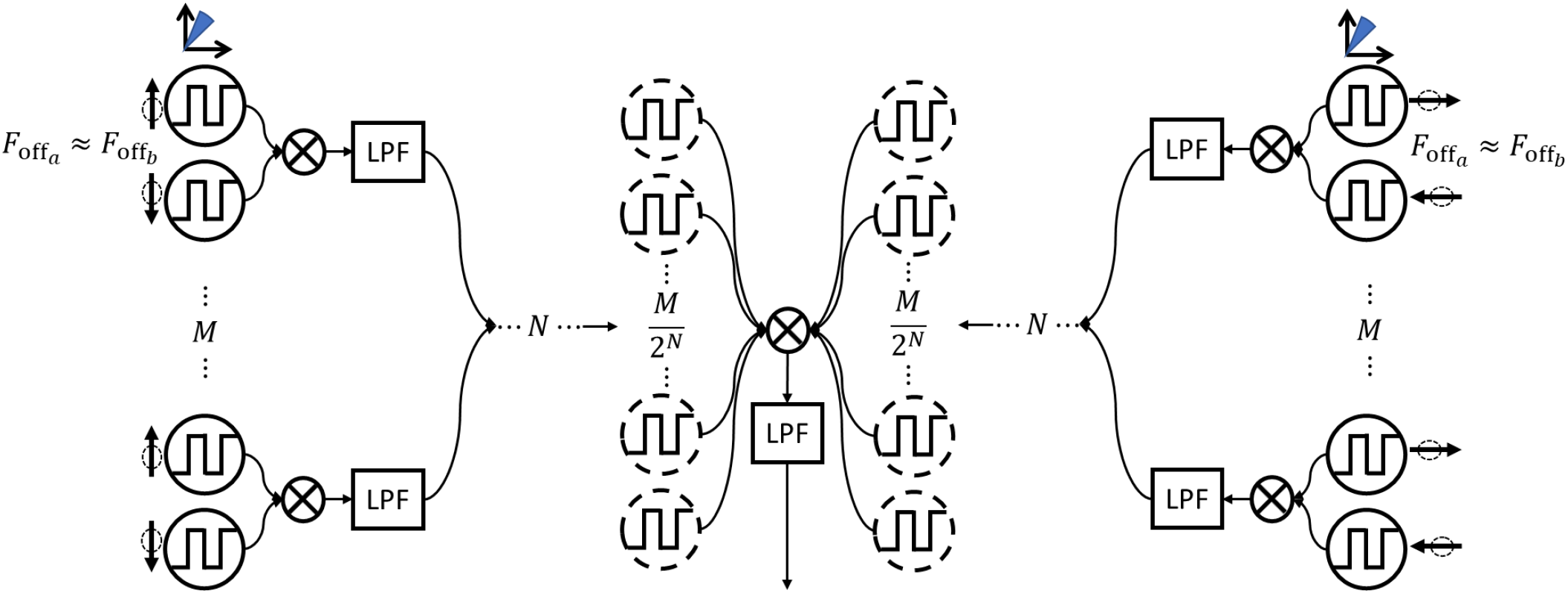
Structural overview of the construction method coupled with offset-reduction to form Place cells with theta oscillators with variations in β and base frequency, i.e. non-zero F_off_.

**Figure 8:**
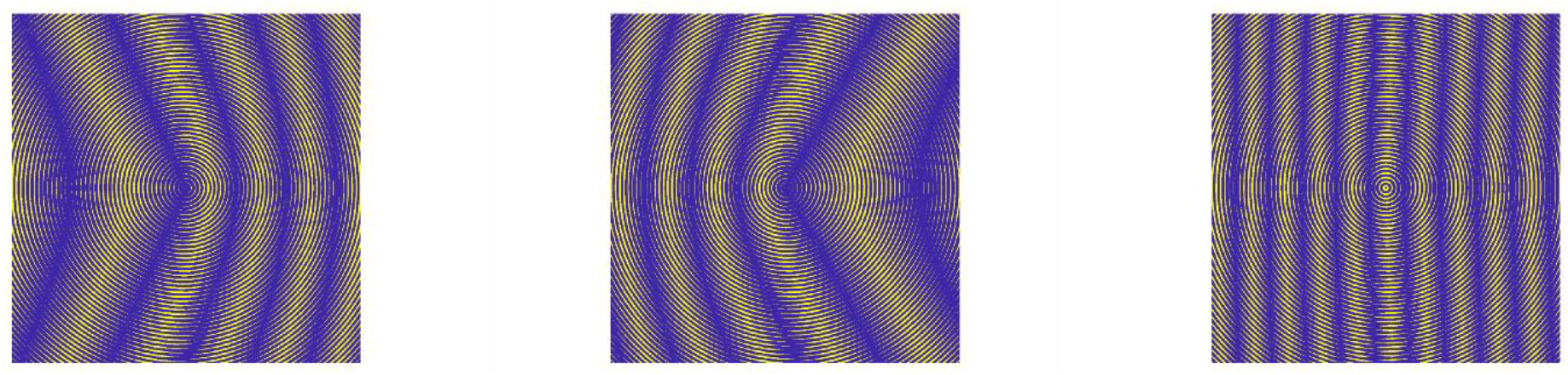
Demonstration of offset-reduction. The left figure is a Theta cell with 0° preferred direction and 0.5 Hz offset frequency interfered with by an idling frequency; the middle one is another with 180° preferred direction and 0.6 Hz offset frequency. The right one is the pattern from interfering the two Theta cells, exhibiting L halved and suppressed offset frequency (i.e., less curvature in the grid)

Though the magnitude of the offset frequency is reduced, the remainder, (*F*_off_*a*_ − *F*_off_*b*_), would still contribute to displacement-independent phase accumulation, thus affecting the Place cell’s spatial encoding accuracy. To mediate this problem, we adjust the phase computation applied to the effective theta oscillator so that it takes the effective offset frequency into consideration, resulting in equation (15).

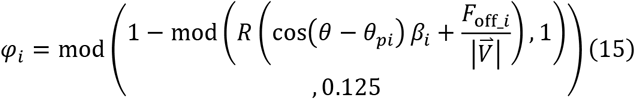

The combination of the offset-reduction and modified phase-shift computation allows us to form Place cells in a way that handles variations in both frequency response factors and idling frequencies. To design our system succinctly, we perform AND operations between all the interference signals. This means that the Theta cell with the largest frequency response factor *β*, and thus the thinnest stripes according to equation (4), will determine the size of the Place field. Due to the complication of the introduced offset-reduction, the size of the Place fields is controlled not through the individual Theta cells, but through the effective *β* oscillator after offset-reduction.

## 6. Path tracking

With the capability of forming a Place cell at any spatial location with realistic performance of Theta cells, it is possible to produce a number of Place cells to tessellate the space for location- and path-tracking. Based on the size of the place field of the Place cells formed and the region of interest, the required density of the Place cells can be assessed. This allows for at least one Place cell to be active during a tracking event, which enables the system to keep track of the subject’s location based on which Place cell fires in an egocentric frame.

As stated before, due to the impact of the remaining offset frequency, the phase equation for a Place cell is dependent on the speed and time involved in traveling from origin to the Place cell’s designated spatial location. This makes the Place cell function only when the subject is traveling with the preset speed and route, such as a straight line as assumed in equation (12). When traveling at a different speed, the Place cells near the origin would still track normally because the time needed to travel to those locations is small, and not much phase is accumulated by the offset frequency. Thus, for the collection of Place cells to function, we imposed a restriction that, during each tracking event, the velocity is held constant.

However, it is natural to change velocity while navigating through the environment, thus, tracking across variation in both speed and direction is crucial for reaching a desired target and path encoding. But the proposed Place cells will only function properly when two conditions are met: velocity is constant, and the tracking does not persist for too long, due to the periodicity of the Place cells. To resolve this conflict, we introduce a phase-reset mechanism for the collection of Place cells. It will pull all the Theta oscillators back to a synchronized zero phase, which is equivalent to clearing the phase accumulation and bringing the tracking back to the origin, setting up a new egocentric frame. Upon this resetting, the latest-fired Place cell will be recorded. The reset will be triggered by two events: if the velocity changes, or if the time since the last reset has reached a preset interval which shall be smaller than the Place cells’ common period. Similar phase resetting phenomena have been observed in Theta rhythm of hippocampal cells, though, however without clear evidence of whether it correlates to a change in velocity as we are proposing here [31]. In both the OI model and CAN model, phase resetting is also introduced to clear off the effect of path integration accumulated error in both the direction and magnitude estimation of the travel velocity [7-8], [10], and could serve the same purpose in this model as well.

Referring to equation (12) for Place cell phase computation, the Place cells’ connections only depend on the speed, due to the nonzero offset frequency term, which means that one collection of Place cells could track travels with a constant speed regardless of direction. But a new collection of Place cells is then necessary for tracking under a different speed. So, in our model, a set of precomputed collections of Place cells is used for tracking. Each collection is differentiated by the speed it accommodates. Upon the phase-reset event, if the velocity changes only in direction, the tracking will still happen within the same collection of Place cells. But if the magnitude changes, then tracking needs to be taken over by another collection of Place cells. Then, the Place cell that gets recorded will implicitly contain the speed information while traversing through this segment. And because we need the same number of Place cells to cover the region, and they operate with the same time interval, each collection operates on a different spatial scale and accuracy as well. This is, potentially, a nice feature: while the speed is slow, the place field is smaller and thus more accurate in tracking and recording, and vice versa. Moreover, this change of scale will also be reflected in the Grid cell model, resulting in the grids having different spatial densities, which provides a potential explanation for parallel multi-scaled Grid cells observed in rodents [25], and further support the models that take the advantage of multi-scaled Grid cells for place cell formation [16][29].

## 7. Results

The performance of the proposed model is now assessed in simulations, assuming Gaussian distribution of both frequency response factor *β* and offset frequency *F*_off_. Though the 60° difference between the two sets of Theta cells (as demonstrated in figure 4) is feasible, we want to emphasize the implementation’s friendliness. Thus, in the performance assessment, we adopt a 90° difference in the preferred vectors to better correspond to the standard Cartesian coordinate system. Two sample Place cells generated with our proposed model with *β* and *F*_off_ generated from their respective normal distributions are shown in figures 9(a, b). We then compute the mean squared error of the spatial pattern between each trial and an ideal Place cell created through Welday et al.’s method, as shown in figure 9(c). For each configuration of specified number of Theta cells and standard deviation of offset frequency *F*_off_, 100 trials were generated and an average mean squared error is plotted in figure 9(d, e).

**Figure 9:**
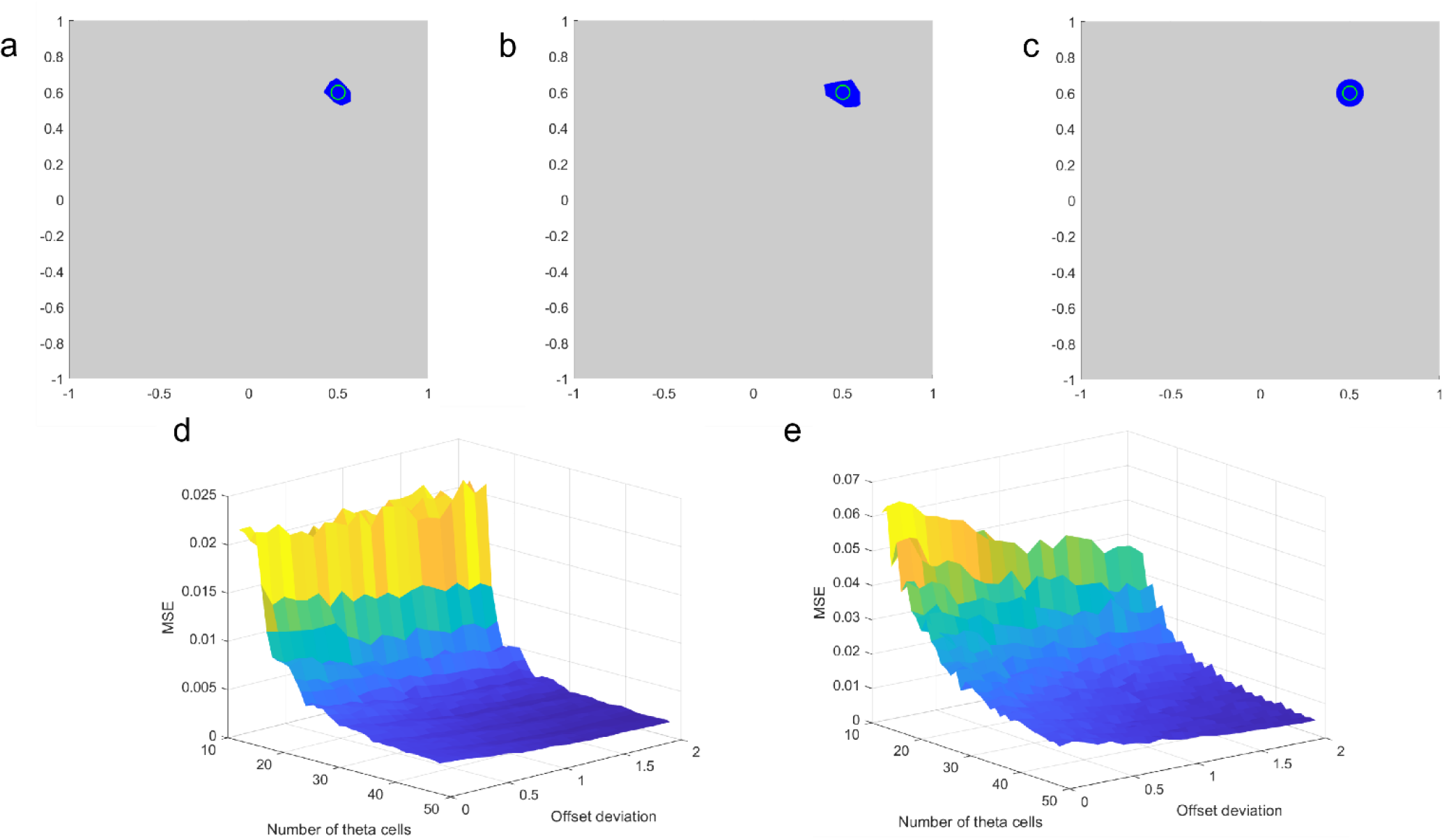
Performance assessment. Figure (a) is a sample Place cell pattern generated from 48 Theta cells with frequency response factors generated from three Gaussians with distinct means, 0.2*[3 5 7]Hz per unit, and a common standard deviation of 0.2Hz per unit, with random idling frequencies with a mean of 8Hz and standard deviation of 2Hz. Figure (b) is a sample Place cell pattern generated from 48 Theta cells with frequency response factors generated from Gaussians with a common mean of 1Hz per unit with standard deviation of 0.2 Hz per unit, and random idling frequencies with a mean of 8Hz and standard deviation of 2Hz. Figure (c) is a baseline ideal Place cell pattern for comparison. Figure (d) is the average MSE computed between trials like figure (a) and figure (c). Figure (e) is the average MSE computed between trials like figure (b) and figure (c).

Two setups of the experiment are shown here. In figure 9(d), we choose 3 mean values (0.2 × [3, 5, 7] Hz per unit), with the same deviation (0.2 Hz per unit) for the *β*_*i*_ to simulate a similar construction setup as in figure 4. We then vary the number of Theta cells from 16 to 48, and the standard deviation of the offset frequencies from 0 to 2 Hz, with a mean frequency of 8 Hz. The results in figure 9(d) show that, with a small number of Theta cells, trials with small deviations of *F*_off_ would have similar errors as large deviations. Yet, due to the uncertainty of the *β*_*i*_, artifacts are likely to occur while forming a single stripe, resulting in several side stripes like the thin stripes in figures 9(d, h), which create error. Increasing the number of Theta cells, a larger population of *β*_*i*_ provides a more diverse spectrum basis that helps the formation of single stripes per direction, similar to the Fourier decomposition of a delta function, thus driving down the error. Moreover, the trials with larger deviations in offset frequencies introduce bending to the gratings, making overlapping at side stripes even less likely, which reduces the error even more with the sample Place cell shown in figure 9(a). On the other hand, the offset-reduction strategy with a larger pool of choices confines the effective offset frequencies to small values that will not alter the directivity of the gratings as seen in figure 6(a). Together, the least amount of error occurs when there is a large deviation in the *F*_off_ with a large number of Theta cells, *suggesting that, under our proposed model, variations among the Theta cells are in fact a contributing factor for accurate Place cell formation*. This may also be consistent with biological observations, where typically a large number of neurons converge to implement any particular functions in cortex, coincides with the Place cell being a pyramidal cell [3][5][20].

This phenomenon can be further illustrated by the second experiment where *β*_*i*_ are generated from a single normal distribution with a mean of 1 Hz per unit, and a standard deviation of 0.2 Hz per unit, shown in figure 9(e). The variations alone form a unique stripe and, further, a Place cell in the space. It produces very similar results when there is a greater number of Theta cells available; *in these cases, trials with high deviations for offset frequency exhibit an advantage over those with lower deviations*. But, unlike the previous setup, when the number of available Theta cells is small, the trials with higher offset frequency deviations also prevail, since the model can only rely on the bending of stripes to avoid the formation of side stripes or dots when the diversity of *β* values is restricted. This restriction continues to affect the accuracy even when the number of Theta cells increases, resulted in a larger place field in figure 9(b), and also a higher overall MSE than the previous setup. This result further demonstrates that the combined effect of diversity in both frequency response factors and offset frequencies contribute to a unique Place field for the Place cell in our model.

To assess localization, we first need to populate the area of interest with Place cells. We use the same Theta cell setup from the first experiment—48 Theta cells and a 2 Hz standard deviation—to conduct the localization and path tracking experiment. Based on the size of the place field, as shown in figure 9(a), we can determine the density of Place cells to cover the region. In the following simulations, we generate a collection of 21×21 grid of Place cells to tessellate a 2D space where coordinates *x* and *y* range from −1 to 1, so at least one Place cell is available to fire at any given location. Such coverage enables the system to keep track of the location of the subject based on which Place cell fires in an egocentric frame.

A localization event is demonstrated in figure 10, where the Place cells assigned to the locations along the travel direction fire sequentially while the subject moves at a constant velocity. If the subject changes velocity, the reset event is triggered, which includes recording which Place cell last fired. Each Place cell that gets recorded coincides with the mathematical concept of a vector where a Place cell encodes its terminal point. Upon each phase-resetting event, the Place cells form a new egocentric field and record its relationship with the previous field much as a transformation matrix computes frame relations in robotics, as shown in figure 10(d, h, l, j). A path can be retrieved by sequentially going through those recorded Place cells, just like vector addition. Yet the Place cells also have a third dimension to record the speed for each segment, since the offset frequency forces us to use a multi-scale Place cell organization that implicitly has a dynamic spatial resolution and speed encoding as discussed in section 6.

**Figure 10:**
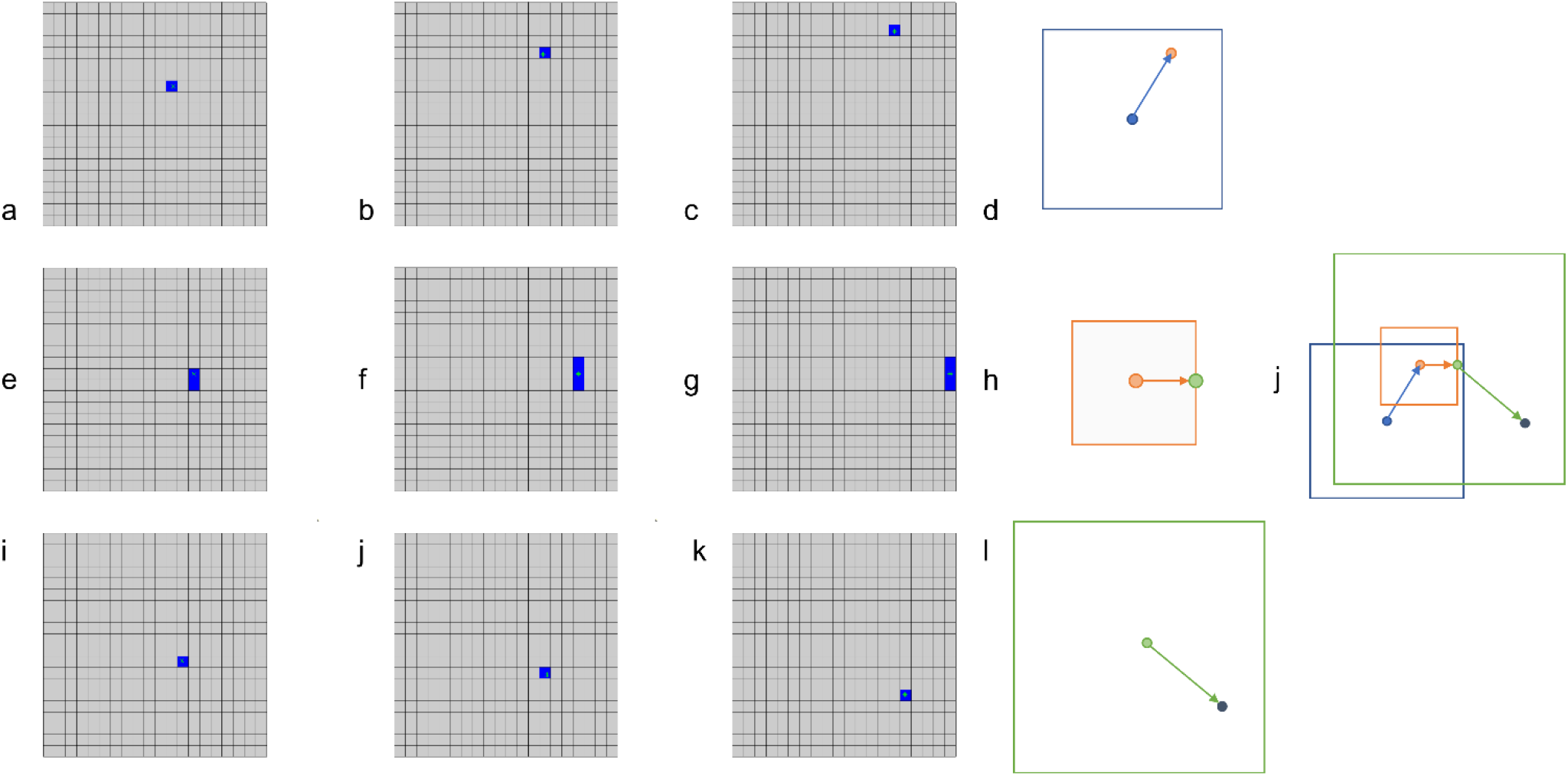
Place cells with designated spatial locations along the direction of travel fire sequentially as the subject travels. A tracking event with 3 speed segments consecutively is simulated. Figures on the left are population maps of 21 by 21 place cells’ instant firing status. Each row of 3 maps on the left shows the sequential firing of place cells while the subject is moving with distinctive constant speeds (i.e, (0.5, 1, 1.5)) and direction. Each row’s Place cell population is designated for that speed only thus has different spatial scale as shown in the conceptual figure in d, h and l. A phase reset is triggered when a change of speed occurs at the time instance shown in c, g or k, which brings the tracking to the center/origin of the next row’s population of Place cells, representing a different measure of space. If the Place cells fired in c, g and k are recorded, a path is recorded in a vector-like fashion as in j along with each segments’ speed and scale information.

## 8. Discussion

In this paper, we first present a oscillatory interference model for constructing Place cells through a set of Theta cells with mutually prime frequency response factors and two distinct preferred directions. We then improved the model to accommodate the possible variations of idling frequencies and frequency response factors among the population of Theta cells. We proposed a novel offset-reduction strategy for interference, and a modified equation (15) to accomandate remaining offset frequencies and variations of frequency response factors. Through simulations, we found that the proposed model benefits from the variations of frequency response factors among Theta cells for forming a unique place field. Moreover, we proposed a subsequent localization and path-tracking mechanism, incorporatingphase resets, to correct the deviation and drift caused by the variations in idling frequencies and sensory errors.

When experimenting with the model, we adopt square-wave oscillation and quantized phases to demonstrate the model’s suitability for hardware implementation. Our team built a mixed-mode theta chip [32] to imitate the behavior of Theta cells described in equation (1), which has substantial variations across all Theta cells, as figure 11 shows. With the proposed model, variations might be a beneficial feature for construction of Place cells.

**Figure 11:**
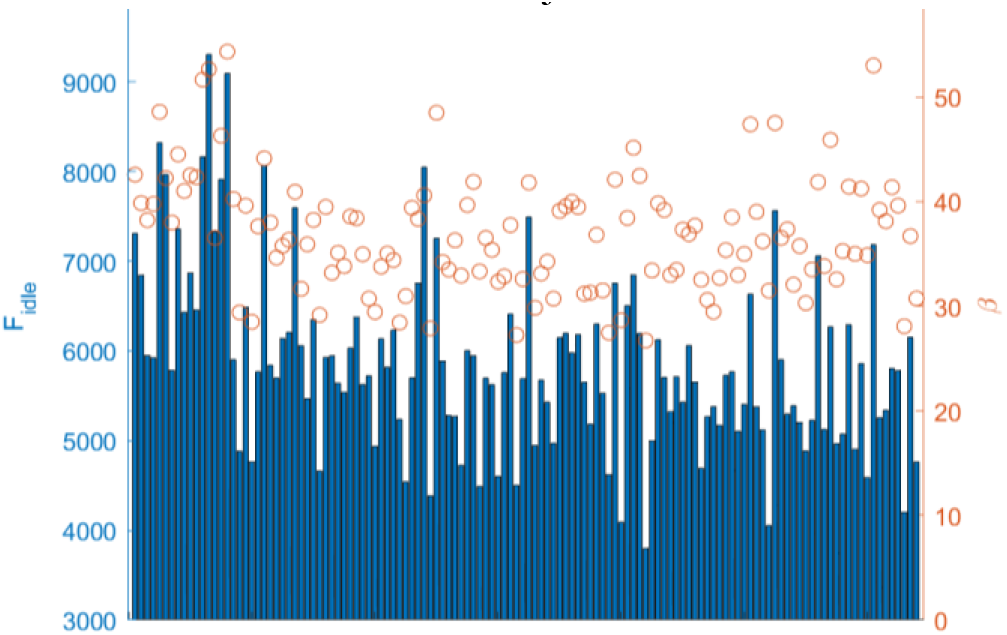
Theta Cell chip’s 128 oscillator units parameterized response after fitting with the Theta cell equation (to be published). Bars showing idling frequencies and red dots for frequency response factors.

To better incorporate the square wave oscillation, our model utilizes a multiplicative interference rather than additive. In terms of biology, it resembles the conductance inhibition that has been observed in the hippocampus CA3 [28]. In terms of computation complexity, it can be more simply implemented with a combination logic AND operation in hardware, compared to additive interference which requires two stages of operation and a memory unit. As an example, a place cell form through Welday et al.’s model using 12 ideal theta cells, i.e. perfectly matched in frequency and gain, needs 11 additions and 1 comparison to compute the status of place cell in a time slice, whereas our model requires 47 Boolean AND operations from 48 non-ideal theta cells that have variations. As a reference, a typical cell in the continuous attractor model requires a weighted inhibition from all the neighboring cells within a certain radius of closeness. Even for a small radius of 4 surrounding layers of cells, 64 multiplications and additions needed to be computed for one timestep for that cell only.

Throughout our model, as many have, we hypothesize a purpose for the Grid cells. In figure 4, we demonstrate that Grid cells could be formed as a middle step for generating Place cells. Combined with our offset-reduction mechanism, Grid cells might have a further purpose of pairing two Theta cells with closely matched base frequencies together to suppress its impact on the Grid cell behavior just as we did. The only difference is that, instead of pairing two Theta cells with preferred directions 180° apart, Grid cells pair those with 60° or 120° differences. A construction rule for forming the Grid cell would then just entail reinforcing synapses from the Theta cells with the closest frequencies.

One development of the current model that we plan to investigate is to reduce the connections involved in forming a Place cell. As shown in figure 9, it seems that, the more Theta cells, the better, when we have a fully connected interference network forming each Place cell from all available theta-cell pairs. However, we would like to develop a standard to identify each theta-cell pair’s contribution to the Place cell, and lesion/prune those that have negative or neutral impact on forming the Place cells, so as to reduce the network complexity. We would also implement the model in hardware using our Theta Cell chip, to further validate the model. The latter will be the subject of a future submission.

## Acknowledgements

The authors would like to thank Akwasi Akwaboah for providing computational resources.

## Notes

### Competing Interest Statement

The authors have declared no competing interest.

